# Skuas as sentinels of high pathogenicity avian influenza H5N1 on the Antarctic Peninsula in the 2024/2025 austral summer

**DOI:** 10.64898/2026.02.15.706047

**Authors:** Michelle Wille, Will Abbott, Darrel Day, Yi-Mo Deng, Xiaomin Dong, Ted Gibson, Ryan Hope-Inglis, Michelle McCulley, Ida Olsson, Arvind Varsani, Teri Visentin, Matthew Walters, Meagan Dewar

**Affiliations:** WHO Collaborating Centre for Reference and Research, at the Peter Doherty Institute for Infection and Immunity, Melbourne, Victoria, Australia; Department of Microbiology and Immunology, University of Melbourne, at the Peter Doherty Institute for Infection and Immunity, Melbourne, Victoria, Australia; Intrepid Travel, Sydney Australia; Consulting Antarctica, Sydney, Australia; Division of Health, University of Waikato, Private Bag 3105, Hamilton 3240, New Zealand; Biodesign Center for Fundamental and Applied Microbiomics, Center for Evolution and Medicine and School of Life Sciences, Arizona State University, Tempe, USA; School of Life and Environmental Sciences, Deakin University, Geelong, Australia; School of Biological Sciences, University of Canterbury, Christchurch, NZ; Future Regions Research Centre, Federation University of Australia, Berwick, Victoria, Australia

**Keywords:** Antarctica, avian influenza virus, disease ecology, high pathogenicity avian influenza, HPAI, H5N1, Skuas, Penguins

## Abstract

Despite Antarctica’s geographic isolation, the first incursion of high pathogenicity avian influenza (HPAI) H5N1 was detected in the 2023/24 austral summer. Surveillance for HPAI H5N1 in Antarctica remains patchy due to logistical, financial, and infrastructure challenges, with many suspected cases remaining unconfirmed, and few viral genomes sequences available to date. Through the 2024/25 austral summer we undertook five sampling expeditions to the South Shetland Islands and Antarctic Peninsula facilitated by cruise ships/operators. Across more than 500 faecal environmental samples collected from apparently healthy penguins and marine mammals, we found no detectable evidence of HPAI H5N1. However, HPAI H5N1 was detected in all but one of the skua carcasses sampled, which, in most cases, were found within meters of penguin sub-colonies. All HPAI H5N1 viral genomes sequences from skuas on the Antarctic Peninsula fell within a single lineage, which included those genomes from skuas sampled in the 2024/25 season from the South Shetland Islands. Genomes were in a different clade to those from the Antarctic Peninsula collected in the 2023/24 austral summer. Our results confirm although the prevalence may be low, HPAI H5N1 is established in Antarctica, emphasizing the need for ongoing surveillance to monitor and mitigate threats to wildlife, even in the planet’s most isolated regions.

## Introduction

High pathogenicity avian influenza (HPAI) H5N1is causing a global panzootic, and since 2021 has spread to every continent except Australia [1,2]. It has been catastrophic for wildlife, with detections in over 400 wild bird and 60 wild mammalian species [3,4]. Beyond a broad hsot range, HPAi H5N1 has had detrimental population and species level consequences for some species. For example, 40% of Peruvian Pelicans (*Pelecanus thagus*) in Peru died in 2022, 60% of all Northern Gannets (*Morus bassanus*) across the North Atlantic died in 2022, and 10% of all South American Sea Lions (*Otaria flavescens*) in South America died in 2022-2023 [5–7]. Mass mortality events at this scale not only have species level impacts, but will certainly also have ecosystem-level impacts, the extent of which will likely only be revealed in the years to come.

Prior to 2023, the remote continent of Antarctica and associated fauna were considered beyond the reach of HPAI H5N1 outbreaks. However, Antarctica’s interconnectedness with other regions, and particularly South America are increasingly clear, with the emergence of HPAI H5N1 in the sub-Antarctic and Antarctica in the 2023/24 austral summer. HPAI H5N1 arrived in the sub-Antarctic island of South Georgia in October 2023 [8], and was first confirmed on the Antarctic Peninsula in February 2024 [9]. Over the course of the 2023/24 austral summer, HPAI H5N1 was detected across much of the Antarctic Peninsula, with the most southern detection occurring in Marguerite Bay [10]. HPAI H5N1 resurged in spring 2024, and was reported along the Antarctic Peninsula throughout the 2024/2025 austral summer [10]. In addition to active circulation in fauna on South Georgia, the Falkland (Malvinas) Islands, and the Antarctic Peninsula, HPAI H5N1 spread dramatically through the sub-Antarctic, with detections on the sub-Antarctic Islands of Gough Island, Marion Island, Crozet Island, Kerguelen Island and Heard Island [10–13]. Since its arrival on the Antarctic continent, it has been detected in diverse avian (n = 11) and mammalian (n = 5) species in the region. While impacts on the birds and seals of the sub-Antarctic islands have been considerable [*e*.*g*. 11,14–16], impacts on the charismatic wildlife of the Antarctic continent have been considerably less than anticipated.

Surveillance for HPAI H5N1 in Antarctica poses a number of unique challenges, mostly attributed to the lack of response teams, limited opportunities for repeated visits, and lack of molecular laboratory facilitates [17]. Therefore, surveillance in the region is patchy, with both confirmed and suspected cases reported by scientists, citizen scientists and tourist operators [10]. Herein we report the outcomes of four months of surveillance activities undertaken for HPAI H5N1 undertaken in the 2024/25 austral summer, in the South Shetland Islands and Antarctic Peninsula onboard two tourist vessels: the MS Ocean Endeavour and MV Argus. This comprises the most comprehensive monitoring study undertaken in the 2024/25 season. We aimed to reveal whether HPAI H5N1 was present in colonies of apparently healthy penguins, continue to monitor the impacts of this virus on Skuas, and reveal virus diversity in the region.

## Methods

### Ethics statement

All animal experiments were approved by the Institutional Animal Care and Use Committee at Federation University Australia (AE/2003/004). All research was undertaken in accordance with Australian Antarctic Division permit #ATEP 24/4877.

### Sample collection

Five sample collection expeditions were undertaken between November 2024-February 2025. Sample collection was facilitated by Intrepid Tours, on the MS Ocean Endeavour vessel, as well as Consulting Antarctica and EVOKN Expedition, on the M/V Argus vessel (Table S1). Samples were collected at 20 sites situated located on the South Shetland Islands, and the Antarctic Peninsula ranging from the Trinity Peninsula through the Gerlache Strain (Figure 1).

**Figure 1.**
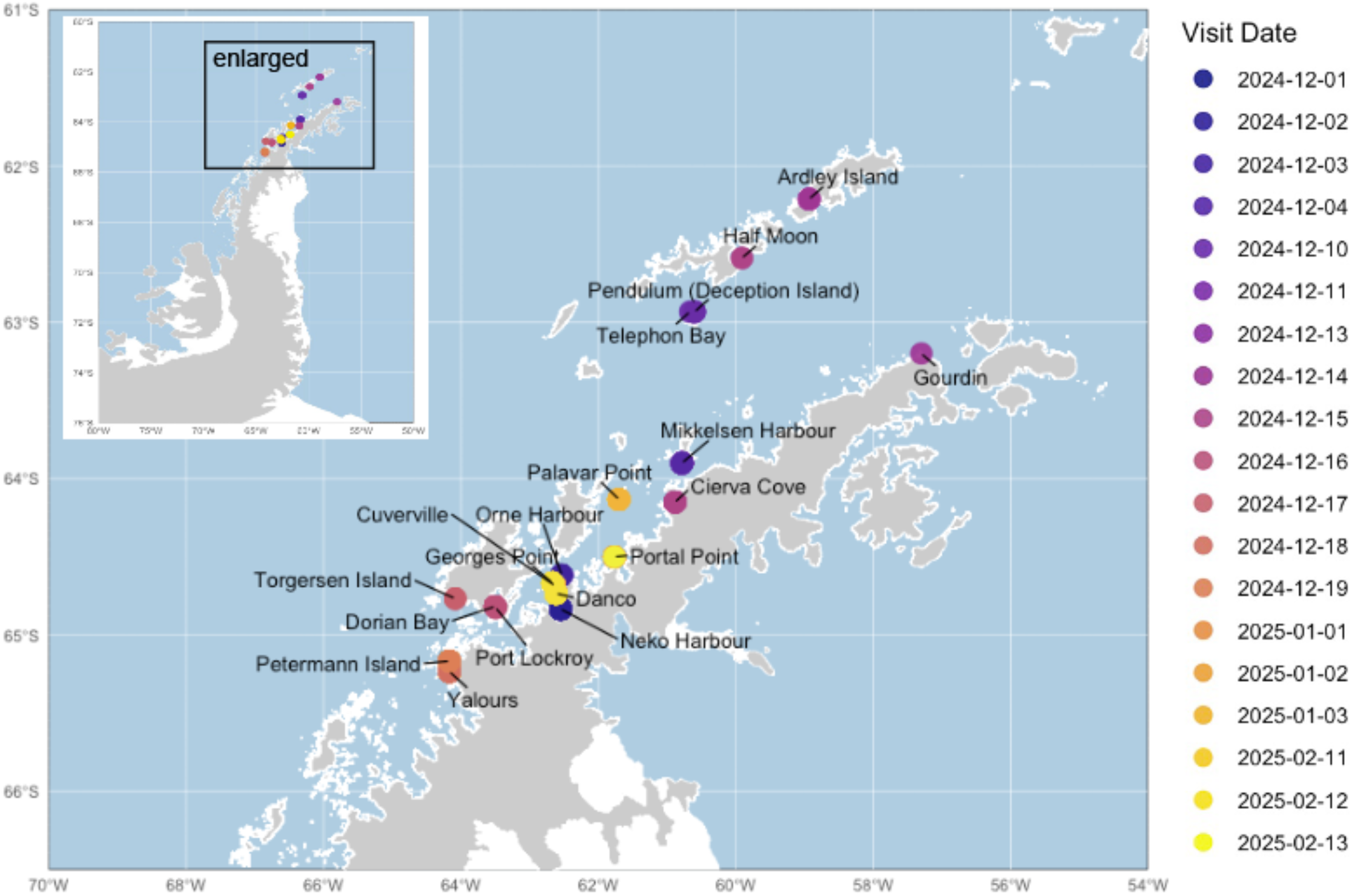
Sampling locations in this study. Points are coloured by the sampling date. Details of sampling locations (including latitude and longitude) for each sampling voyage are provided in Table S1.

Non-invasive faecal environmental samples were collected at penguin colonies. To ensure faecal swab were fresh, penguins were observed defecating, and a swab (Copan FloqSwab) of the faeces was collected. Where possible, swabs of fresh pinniped faeces were collected from zodiac boat landing sites and ice floes. From any carcasses found, we collected oropharyngeal, cloacal and/or brain swabs were collected where possible. Swabs were placed into 1mL of DNA/RNA Shield (Zymo Research) and stored at room temperature on the cruise ship (in alignment with manufacturers recommendations). Samples were placed in −80 °C upon arrival in the laboratory (generally within 2 weeks to one month of collection).

### Sample screening

RNA was extracted from samples using the Qiagen QIAmp 96 Virus QIAcube HT Kit (Qiagen) on a QIAextrator robot. We confirmed that the chemistry of this kit was compatible with DNA/RNA Shield through a comparison of isolates spiked into both virus transport media and DNA/RNA Shield (Zymo Research), with no significant difference in Ct values found. In cases where samples had substantial amount of faecal material preventing the vacuum from removing all the liquid from the columns during the extraction, they were re-extracted using the Quick-RNA MagBead Kit (Zymo Research) on a KingFisher Apex Instrument (ThermoFisher). All positive samples were also re-extracted using the Quick-RNA MagBead Kit (Zymo Research).

Extracted RNA was assayed for a short fragment of the conserved matrix gene [18] of influenza A virus using the SensiFAST Probe Lo-Rox qPCR Kit (Bioline Meridian Science) as previously reported [19]. Positive samples were subsequently tested using the CDC A/H5 (Asian Lineage) subtyping Panel (VER 4) and SensiFAST Probe Lo-Rox qPCR Kit (Bioline Meridian Science), with Ct value cut-offs as per the recommended assay guidelines. Positive samples were also tested using a 2.3.4.4b pathotyping assay [20] with the Qiagen One Step RT-PCR kit and using mastermix and thermocycling conditions described in [20]. Both low pathogenicity avian influenza (A/Turnstone/King Island/7164CP/2014(H6N8)) and HPAI H5N1 (A/Black-faced Spoonbill/Hong Kong/AFCD-HKU-22-21944-01009/2022(H5N1)) RNA were used as controls for all assays.

### Hybridization capture sequencing

Initial amplicon sequencing [undertaken as per 21] was not successful, such that we leveraged a hybridisation capture sequencing approach. Samples with a Ct value of less than 31 were selected for sequencing, with one sample per carcass attempted (the sample type with the lowest Ct value). Of these, we included ARG071 and ARG072 (reported as A/Brown skua/Torgersen Island/o8182/2024) as positive sequencing controls; they were sequenced in McCulley *et al*. (2025) on board the MV Argus within 24hrs of collection [22]. These two HPAI H5N1 virus genomes have not been included in subsequent analysis as they were used as control samples and have been previously reported.

Hybridisation capture sequencing was performed using a custom panel (Twist Bioscience, TE-92404697, Table S2), with the Twist Total Nucleic Acids Library Preparation EF Kit 2.0 (Twist Bioscience) for Viral Pathogen Detection and Characterization. The custom probe panel targets seasonal and zoonotic influenza, respiratory syncytial viruses, and human metapneumovirus (Table S2 for the list of influenza viruses used in the probe panel design). The hybridisation capture sequencing was based on Twist Target Enrichment Standard Hybridization v1 Protocol with modifications listed here: all RNA samples were treated with ezDNase (Invitrogen, ThermoFisher Scientific) to reduce host genome DNA contamination before double-stranded cDNA synthesis; a reduced volume of one quarter of reagents was used for double stranded cDNA synthesis, metagenomic library preparation and target enrichment; DNA fragmentation was done at 37 °C for 10 minutes; Twist UMI Adapters for adapter ligation were used for library preparation. Eight metagenomic libraries were multiplexed into one hybridization capture reaction and resulting enriched libraries were sequenced on an Illumina iSeq with 150bp paired end reads. Demultiplexing and quality control of reads, and genome assembly was undertaken using the IRMA (Iterative Refinement Meta-Assembler) pipeline (Shepard et al. 2016).

### Phylogenetic analysis

All HPAI H5N1 genomes from Antarctica, the sub-Antarctic region, as well as the Falkland/Malvinas Islands as well as the top BLAST hits for all the genomes we generated were downloaded from GISAID (Global Initiative on Sharing All Influenza Data, https://gisaid.org/) (11/09/2025). The coding regions of all segments were ordered by descending length (from segment 1 to segment 8), and concatenated. While all genomes included contained all eight segments, some segments were only partial (*e*.*g*. genomes from Kerguelen had incomplete PB1 sequences). Sequences were aligned using MAFFT [23] and the E-ins-L algorithm to account for the gaps generated by the incomplete segments, integrated within Geneious Prime (Dogmatics). Maximum likelihood phylogenetic trees incorporating the best-fit model of nucleotide substitution were estimated using IQ-Tree [24] with 1000 ultrafast bootstraps on the University of Melbourne (Australia) high performance computing (HPC) server, Spartan.

The alignment was subsequently filtered to include only genomes for which all segments were complete. We further filtered out genomes comprising different sample types from the same individuals, preferentially retaining either brain or oropharyngeal samples. We evaluated the extent of molecular clock-like structure in the data by performing linear regressions of root-to tip distances against year of sampling using maximum likelihood trees using TempEst [25]. As evidence for a molecular clock was obtained, time scaled phylogenetic trees were estimated using BEAST v1.10.4 [26], under the uncorrelated lognormal relaxed clock [27] and an HKY+G to accommodate for overlapping reading frames in the concatenated genomes. The Bayesian skyline coalescent tree prior was used as this likely reflects the complex epidemiology dynamics of avian influenza viruses through time [28]. The trees were run for 100 million generations on the University of Melbourne HPC, Spartan. Convergence was assessed using Tracer v1.8 (http://tree.bio.ed.ac.uk/software/tracer/). Maximum credibility lineage trees were generated using TreeAnnotator following the removal of 10% burnin, and trees were visualised using Fig Tree v1.4 (http://tree.bio.ed.ac.uk/software/figtree/).

### Mammalian adaptation

To understand whether there was any evidence of mammalian adaptation in the HPAI H5N1 genomes, we scanned genomes for mutations that emerged in South American pinnipeds, listed in [29]. We further interrogated lists of significant mutations generated by FluServer (http://flusurver.bii.a-star.edu.sg), using the “automatic detection of closest reference (among current vaccine strains, full genomes)” option, as well as using A/Baikal teal/KoreaDonglim/3/2014(H5N8) as the reference.

### Analysis and plotting

All analysis was undertaken using R 4.3.3.3 integrated into R Studio 2025.05.1+513 (2025.05.1+513). Maps were made using ggplot2, with the base map from https://data.bas.ac.uk/items/4ecd795d-e038-412f-b430-251b33fc880e/. Ct values were compared using linear mixed model with carcass ID as a random effect using the lme4 package [30]. We restricted the analysis to carcasses from which at least two different sample types were collected.

## Results

### Faecal sampling of apparently healthy animals

Faecal samples (n=523) were collected between November 2024 – February 2025 across 5 expeditions (Table 1, Table S1). Most samples were collected from the Antarctic Peninsula (n=507), and in particular the Gerlache Strait. A further 16 samples were collected from the South Shetland Islands (Figure 1). Most faecal samples were collected from penguin species including Adélie Penguin (*Pygoscelis adeliae*; n=76), Chinstrap Penguin (*Pygoscelis antarcticus*, n=31), and Gentoo Penguin (*Pygoscelis papua*; n=340), as well as other avian species in small numbers (Table 1). Faecal environmental samples were also collected from a small number of pinnipeds, including Weddell Seals (*Leptonychotes weddellii*; n=15), Antarctic Fur Seals (*Arctocephalus gazella*; n=3), Southern Elephant Seals (*Mirounga leonina*; n=1), and unidentified seal species (n=2). All samples were negative for influenza A virus.

**Table 1:**
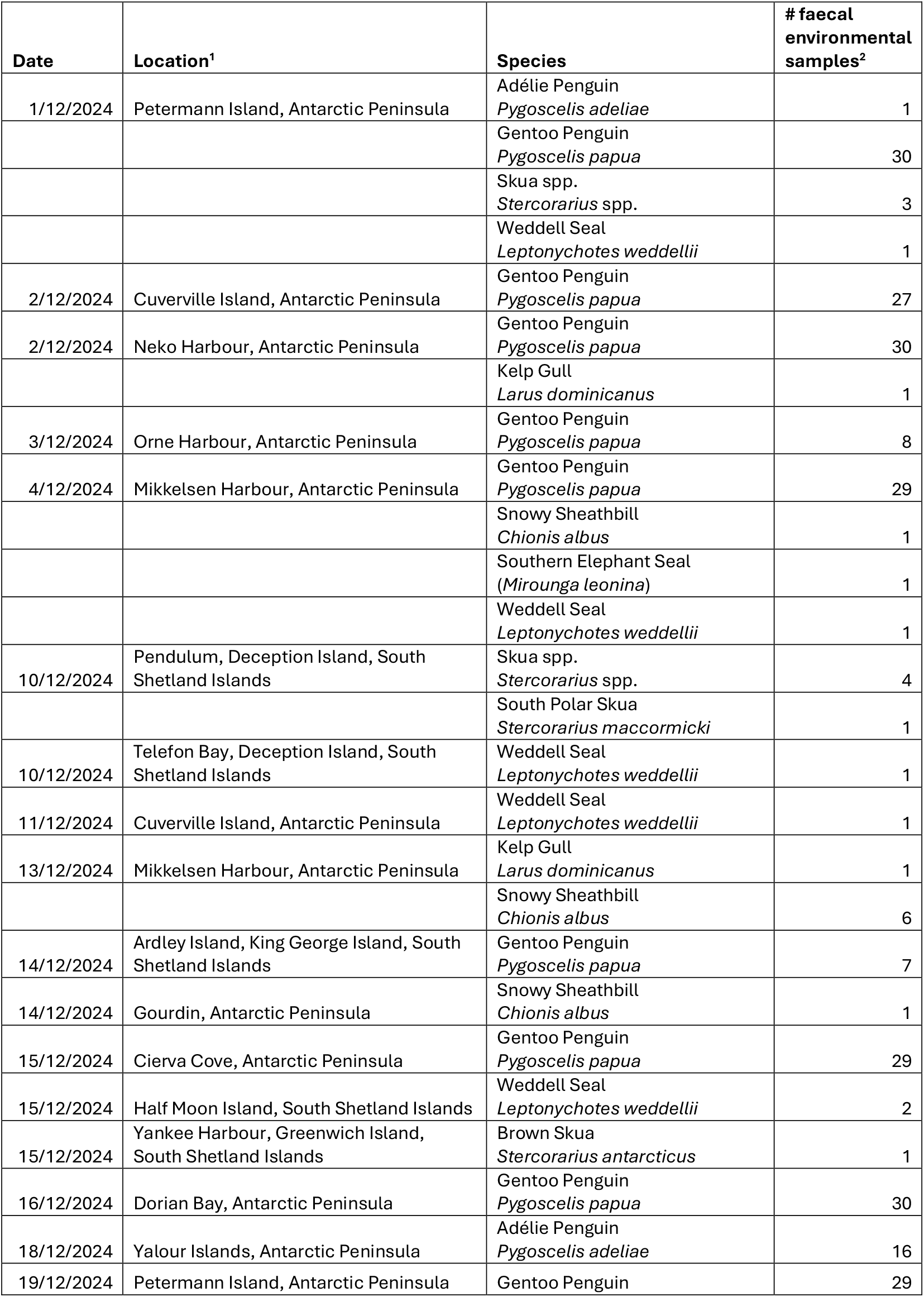
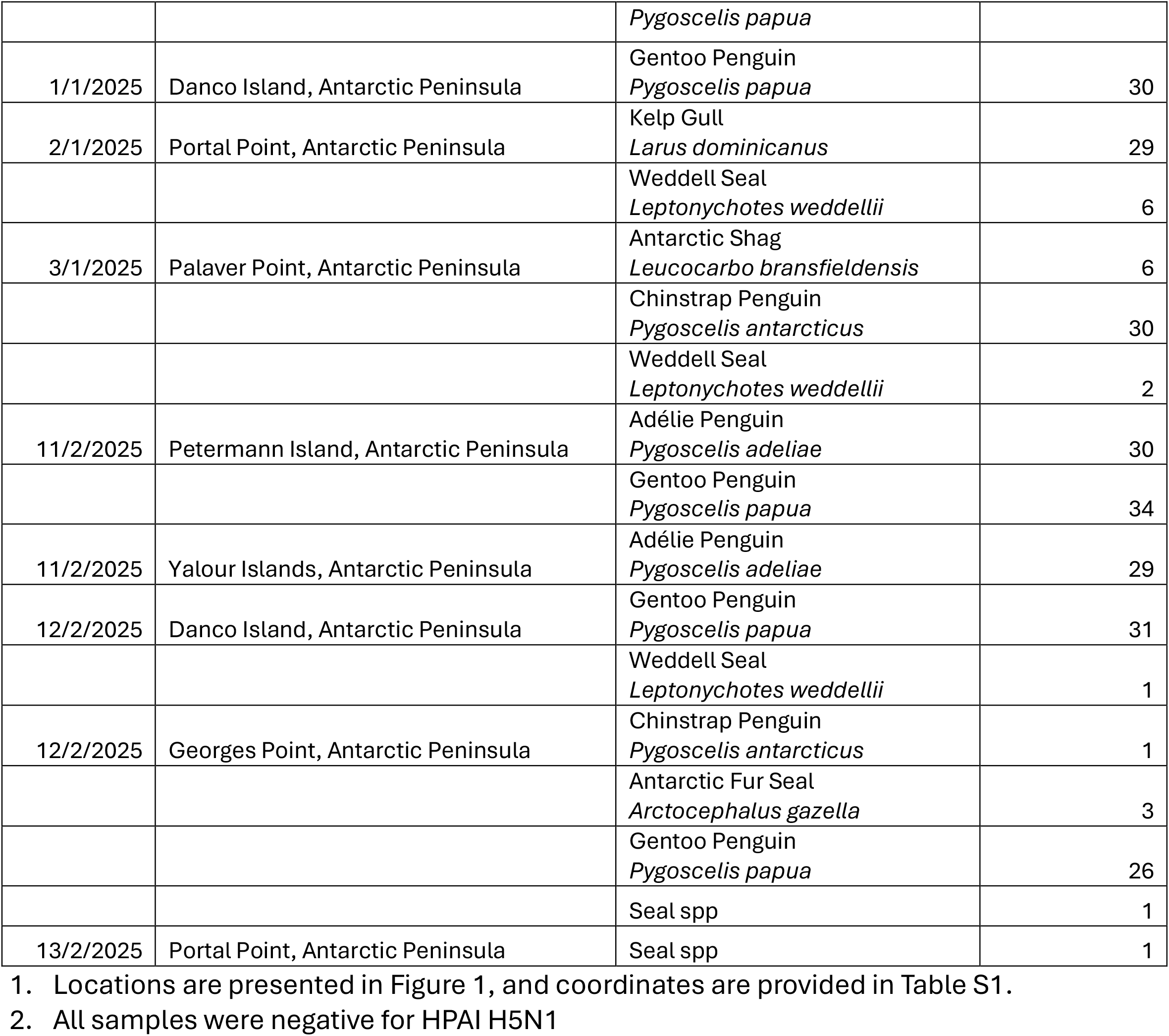
Faecal environmental samples from apparently healthy animals collected in this study.

### Overrepresentation of HPAI detections in Skuas carcasses

Thirty-four samples (including oropharyngeal, cloacal and/or brain swabs) were collected from 15 carcasses, of which 14 were from Skua species (*Stercorarius* spp.) (Table 2). HPAI H5N1 was confirmed using a combination of qPCR, and where possible sequencing, in all samples which were positive for influenza A virus. All faecal samples collected from apparently healthy skuas (n=10) were negative. For many carcasses, multiple sample types were collected, including brain, oropharyngeal and/or cloacal. Twelve skua carcasses were positive in at least one sample type; two skua carcasses were negative in all sample types. Sample type had a significant effect on Ct values (F_2_,_17_ = 6.72, p < 0.05), with cloacal and oropharyngeal swabs having higher Ct values compared to the brain swabs (Figure 2). In addition to skua carcasses, we also collected brain, oral and anal samples from a single Weddell Seal carcass, sampled at Port Lockroy on 16/12/2024, which was negative for influenza A virus in all sample types.

**Table 2.** Faecal and carcass samples collected from Skua sp.

| Species | Date | Location | Sample source | If carcass, samples collected? | Result<sup>1</sup> |
| --- | --- | --- | --- | --- | --- |
| Skua spp. | 1/12/2024 | Petermann Island | Faeces |  | Negative |
| Skua spp. | 1/12/2024 | Petermann Island | Faeces |  | Negative |
| Skua spp. | 1/12/2024 | Petermann Island | Faeces |  | Negative |
| Skua spp. | 2/12/2024 | Cuverville Island | Carcass | Oral swab, Cloacal swab | Positive |
| Skua spp. | 2/12/2024 | Cuverville Island | Carcass | Oral swab, Cloacal swab | Negative |
| Skua spp. | 2/12/2024 | Cuverville Island | Faeces |  | Negative |
| Skua spp. | 10/12/2024 | Deception Island | Faeces |  | Negative |
| Skua spp. | 10/12/2024 | Deception Island | Faeces |  | Negative |
| Skua spp. | 10/12/2024 | Deception Island | Faeces |  | Negative |
| Skua spp. | 10/12/2024 | Deception Island | Faeces |  | Negative |
| South Polar Skua | 10/12/2024 | Deception Island | Faeces |  | Negative |
| Skua spp. | 11/12/2024 | Cuverville Island | Carcass | Oral swab | Positive |
| Skua spp. | 11/12/2024 | Cuverville Island | Carcass | Oral swab, Cloacal swab | Positive |
| Skua spp. | 11/12/2024 | Cuverville Island | Carcass | Oral swab | Positive |
| Skua spp. | 11/12/2024 | Cuverville Island | Carcass | Oral swab | Positive |
| Brown Skua | 15/12/2024 | Yankee Harbour | Faeces |  | Negative |
| South Polar Skua | 17/12/2024 | Torgersen Island | Carcass<sup>2</sup> | Oral swab, Cloacal swab<br>Brain swab, Lung Swab | Positive |
| Skua spp. | 2/1/2025 | Cuverville Island | Carcass | Oral swab, Cloacal swab<br>Brain swab, | Positive |
| Skua spp. | 2/1/2025 | Cuverville Island | Carcass | Oral swab, Brain swab | Positive |
| Skua spp. | 2/1/2025 | Cuverville Island | Carcass | Oral swab, Brain swab | Positive |
| Skua spp. | 2/1/2025 | Cuverville Island | Carcass | Oral swab, Cloacal swab<br>Brain swab, | Positive |
| Skua spp. | 2/1/2025 | Cuverville Island | Carcass | Oral swab, Cloacal swab<br>Brain swab, | Positive |
| Skua spp. | 11/2/2025 | Yalour Islands | Carcass | Oral swab, Brain swab,<br>Tissue swab | Positive |
| Skua spp. | 12/2/2025 | Danco Island | Carcass | Oral swab | Negative |

1. Carcasses were deemed positive if any one sample tested positive
2. Carcass reported in [22]

**Figure 2.**
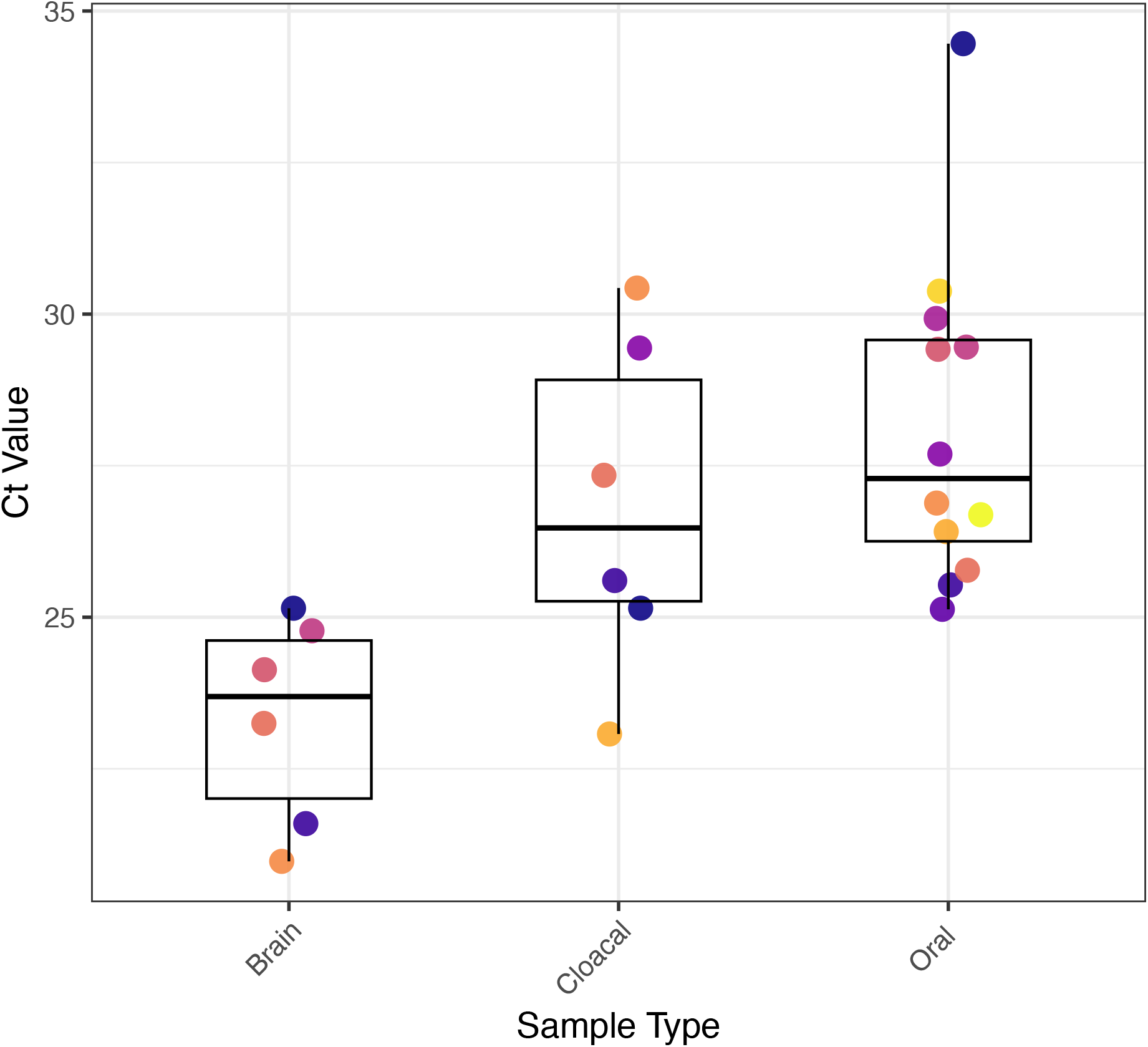
Ct values of different sample types from skua carcasses positive for HPAI. Different colours correspond to different carcasses tested in this study. Only carcasses from which at least 2 swabs (from different sample types) collected are included. Each carcass comprises a unique colour.

One Brown skua (*Stercorarius antarcticus*) carcass sampled on the Yalour Islands was banded, and a query of the banding record indicated the bird was originally banded at Palmer Station in 2001, as a chick, by researchers part of the United States Antarctic Program (USAP). Thus, this individual was 23 years old and was positive for HPAI H5N1. As no other birds were banded, it is unclear whether young or adult birds are being disproportionately impacted.

### Viral genomes from 24/25 are not in the same clade as those from the previous season

We attempted to sequence at least one sample from each positive carcass, with samples selected having Ct range of 23-31. Initial attempts with amplicon sequencing using standard approaches were not successful, however we successfully generated full genome sequences from 15 samples using a custom hybrid capture panel approach. Of the 16 positive samples we attempted to sequence, only one failed (read depth below depth threshold, INT-OE5-452, matrix Ct = 29). For successfully sequenced samples, read depth ranged from a median of 166 (INT-OE5-275, PB1) to 37795 (INT-OE5-281 HA) (Table S3). Both positive control samples (ARG071 and ARG072, which were sequenced in McCulley et al. [22]) were successfully sequenced, validating our use of this sequencing approach. We do not discuss these two HPAI H5N1 genomes further given they have already been reported.

The concatenated HPAI H5N1 genomes from skuas sampled in Cuverville Island (n=12) shared 99.9% pairwise identity to each other, and these genomes clustered together in phylogenetic trees (Figure 3A). Time-structured phylogeny (Figure 3B) suggests these genomes fall into two small sub-clades, where INT-OE5-148, 149 and 150 are sister to a clade containing all the other genomes from Cuverville Island as well as sequences from the South Shetland Islands. Carcasses from Cuverville Island were sampled from December 2024 – February 2025, and sequences from 2024 and 2025 do fall into these different sub-clades. The genome from a skua sampled on the Yalour Islands in February 2025 did not fall into this cluster, rather was sister to the clade containing genomes from Cuverville Island and the South Shetlands (Figure 3). However, the concatenated genome was 99.7% to those from Cuverville Island, and 99.8% similar to that from Torgerson Island reported in [22].

**Figure 3.**
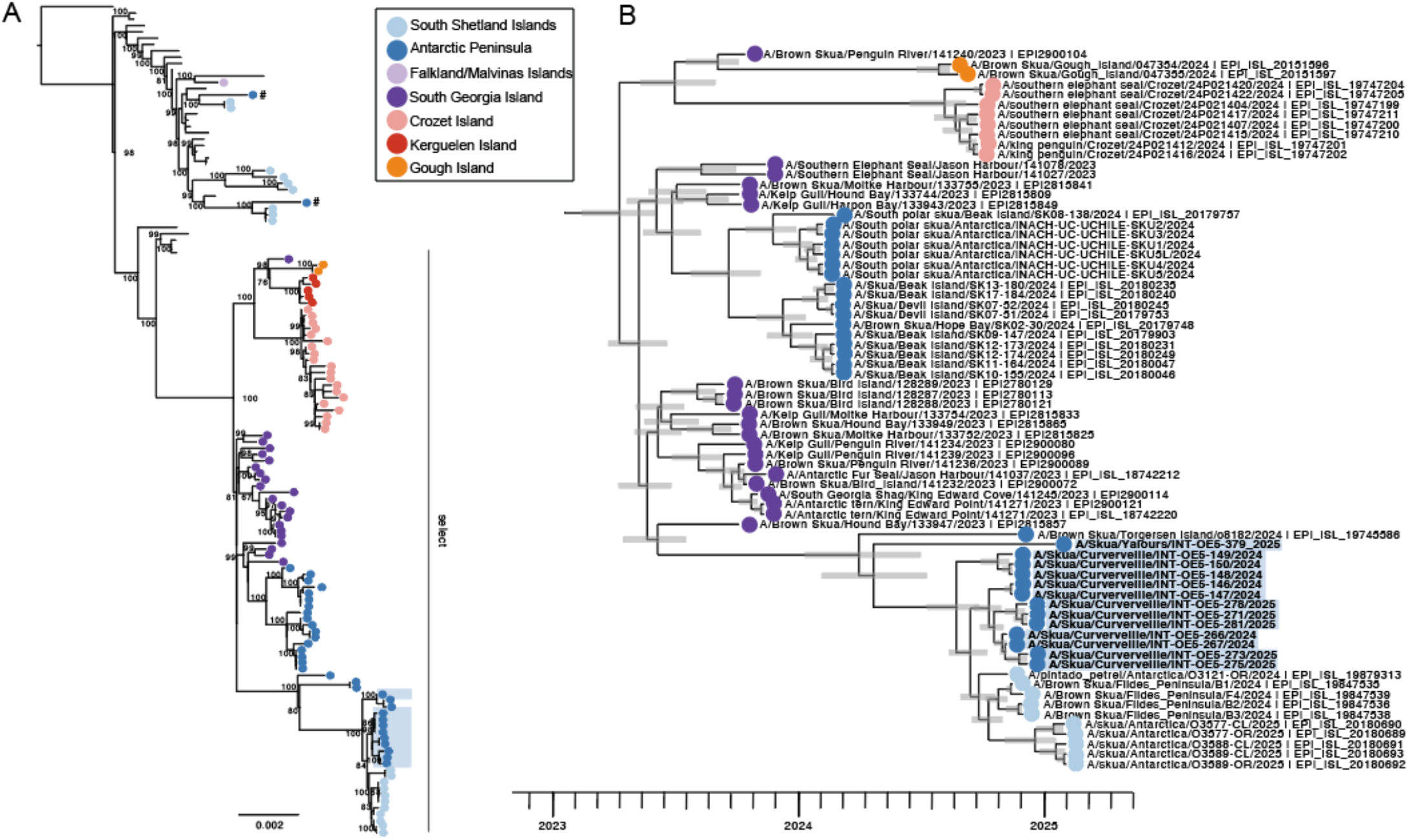
Phylogenetic tree of HPAI H5N1 from the Antarctic Peninsula. (A) Maximum-likelihood tree of the concatenated genomes of HPAI H5N1 in the sub-Antarctic and Antarctica. Bootstrap values are presented at major branches. Scale bar indicates the number of substitutions per site. A/red fox/Michigan (same GenoFlu genotype, B3.2) is used as the outgroup. (B) Time structured tree of the concatenated coding complete genomes of in the sub-Antarctic/Antarctic clade. Time scale is in years. Node bars comprise 95% highest posterior density of node height. Filled circles comprise location. Genomes from this study are indicated by blue boxes. (A, B) Genomes from GISAID downloaded as of 30/11/2025 and for (A) supplemented by from Younger *et al*. (2026) [31], denoted by a hash (#).

Two HPAI H5N1 partial genomes published by Younger et al (2026) [31] reported from the 2024/25 season, from Crab-eater Seals (*Lobodon carcinophaga*) and Kelp Gulls (*Larus dominicanus*), did not fall into the clade comprising genomes generated in this study, but rather were related to genomes from the South Shetland Islands which fell into the broader “South American” clade (Figure 3A). This suggests the potential for further undetected diversity in the region.

Overall, the genomes determined in this study are not in the same clade as those previously reported from the Antarctic peninsula, all of which were from the 2023/2024 season (Figure 3). This suggest that rather than maintenance of the 2023/24 lineage on the peninsula, an independent incursion occurred coinciding with the 2024/25 season, confirming complex patterns of virus movement in the region.

### No evidence for mammalian adapted mutations, but the NA stalk deletion may affect NA glycosylation

No mutations associated with putative mammal-to-mammal transmission in South American pinnipeds [listed in 29], or PB2 E627K, was present in any of the genomes generated in this study. Using FluServer, host specify shift mutations HA T110S and V226A were identified, but only when using H5N8_Baikalteal_2014_KoreaDonglim as a reference, and not when using H5N1_AmericanWigeon_2021_SouthCarolina22-000345-00 (suggesting these mutations are not unique to these viruses). Mutations associated with changes in virulence (PB2 R339K, PB1 T375S, HA S157P, M K101R, and NS L19M, P167S), antigenic drift (NA T65del and S366I, and HA L131Q and S157P), and loss of glycosylation sites (the 51 nt deletion in NA) were identified (NA N58del, N63del, N68del, S70del, T60del, T65del) (Table S3). Many of these mutations in the NA segment were deletions, associated with the 51bp deletion which we and others have observed in some genomes from the Antarctic [22,32,33] (Figure S1).

## Discussion

HPAI H5N1 is having a catastrophic impact on wildlife, globally, with considerable concerns for the unique and charismatic Antarctic wildlife. Indeed, there have been considerable impacts on wildlife in the sub-Antarctic, including the 50% population reduction in the Southern Elephant Seals (*Mirounga leonina*) population in 2023/24 on South Georgia Island [34], approximately 3,000 Black-browed Albatross (*Thalassarche melanophris*) adults found dead at Jason Steeple Island (Falkland Islands) in 2023/24 [35], and more recently a further ~40 dead Black-browed Albatross on Beauchêne Island (Falkland Islands) [36]. However, comparable impacts have not been observed in Antarctica. It is clear that skuas are important indicator species as skua mortality has proven to be the most reliable early indicator of HPAI H5N1 presence [*e*.*g*. 12], and carcasses have tested positive at almost every site tested throughout the Antarctic Peninsula [10]. In addition to Antarctic skua species, Great Skuas (*Stercorarius skua*) in the northern hemisphere have also been heavily impacted [37,38]. Indeed, all positives reported in this study are from skua carcasses. No faecal samples from apparently healthy skuas, or any other species was positive, despite carcasses being found mere meters from penguin colonies. When carcasses were tested for HPAI H5N1, we consistently found that brain samples had the lowest Ct values, which is consistent with neurotropism reported across avian and mammalian species [*e*.*g*.39–42]. Indeed, histology of the H5N1 positive skuas examined during the 2024 Australis Expedition [39], 5/6 had IAV nucleoprotein antigen present in the brain (via immunohistochemistry). Additionally, 2/5 skuas also had IAV nucleoprotein in the lungs, and one was tested positive across a number of other organs [39]. Despite this clear impact on skuas [15,16,43–46], quantitative information on skua population size and breeding success remains scarce for most sites [47–53], which prevents evaluation of whether observed mortality has translated into a population-level decline. Resolving this uncertainty is essential for developing informed conservation and management strategies for these sentinel species.

In the >500 samples from apparently healthy penguins across a diversity of colonies, locations and species, we did not detect HPAI H5N1 in any of these samples. At present, it’s unclear the role that penguins play in HPAI H5N1 ecology. In the sub-Antarctic, HPAI H5N1 has caused mortality events in King Penguins (*Aptenodytes patagonicus*) and Gentoo Penguins [10,11,16]. In South Africa HPAI H5N1 has caused mortality in African Penguins (*Spheniscus demersus*) [54], and in Peru, Humbolt Penguins (*Spheniscus humboldti*) [55], confirming that Sphenisciformes are susceptible, and if exposed, incur considerable mortality. Along the Antarctic Peninsula, it appears that penguin species are not having observed HPAI H5N1 related mortality events given no observations of large numbers of dead penguins or observed colony decline across years in monitored colonies [10]. As to whether penguins are infected with HPAI H5N1 without obvious disease remains unclear, as some studies report HPAI H5N1 in apparently healthy penguins [56], and others do not [46]. Given the uncertainly around the outcome of molecular testing results, serology be critical in clarifying the role of penguins. Indeed, a recent study found no detectable H5 antibodies in any of the Gentoo, Chinstrap, or Adélie Penguins tested [31]. Clarifying whether penguins have subclinical infections is critical. Infected animals with no clinical disease play an important role in virus spread. For example, Mallards (*Anas platyrhynchos*), which have limited disease signs, have been implicated in the long distance spread of HPAI in the northern hemisphere [57,58]. If penguin species have subclinical infections, they may play a similar role in virus spread and maintenance, and should be targeted for far more surveillance than at present. As evidenced by studies in Mallards [*e*.*g*. 59], serology is a powerful approach in helping to resolve viral dynamics in apparently healthy birds, and should be better leveraged in current surveillance.

Given the enormous challenges for HPAI H5N1 surveillance on the continent, viral genome sequences are critical in revealing virus circulation and movement dynamics. The evidence that the H5N1 genomes from the 2024/25 austral summer are not in the same clade as those from 2023/24 suggests that HPAI H5N1 did not persist locally in the environment, such as through overwintering in carcasses or thawed ponds, but was instead reintroduced to the Antarctic Peninsula, most plausibly from South Georgia Island. Limited genomic representation from other regions, including South Georgia Island, prevents firm conclusions about source populations or transmission routes. Overall, this highlights the urgent need for sustained, coordinated genomic surveillance to build a more coherent understanding of HPAI H5N1 emergence and movement across the Antarctic region.

## Data Availability

Genomes were deposited in GISAID (EPI_ISL_20303939, EPI_ISL_20303940,

EPI_ISL_20303941, EPI_ISL_20303945, EPI_ISL_20303946, EPI_ISL_20303947,

EPI_ISL_20304049, EPI_ISL_20304050, EPI_ISL_20304051, EPI_ISL_20304067,

EPI_ISL_20304074, EPI_ISL_20304075, EPI_ISL_20304076).

Phylogenetic tree files and the R script for Figure 1 are available at (https://github.com/michellewille2/Antarctica_24-25/)

## Acknowledgements

We are grateful to the crew of the MV Ocean Endeavour, MV Argus, EVOKN Expeditions, and Brian Kirschenmann for logistical support and facilitating sample collection. We are grateful to Hilda Lau (WHOCCRI) and Matt Neave and Vicky Stevens (ACDP) for assistance in amplicon sequencing attempts.

The CDC Real-Time RT-PCR Influenza Virus A/H5 (Asian Lineage) Subtyping Panel (VER 4) (RUO) (Catalog No. FluRUO-13), FR-1712, was obtained through the International Reagent Resource, Influenza Division, WHO Collaborating Center for Surveillance, Epidemiology and Control of Influenza, Centers for Disease Control and Prevention, Atlanta, GA, USA.

## SUPPLEMENTAL FIGURES AND TABLES

**Table S1.** Sampling expeditions.

| Expedition | Date | Location | Location | Latitude | Longitude |
| --- | --- | --- | --- | --- | --- |
| OE5-03 | 1/12/2024 | Antarctic Peninsula | Petermann Island | -65.16667 | -64.16667 |
| OE5-03 | 2/12/2024 | Antarctic Peninsula | Cuvertville Island | -64.68333 | -62.63333 |
| OE5-03 | 2/12/2024 | Antarctic Peninsula | Neko Harbour | -64.83333 | -62.55 |
| OE5-03 | 3/12/2024 | Antarctic Peninsula | Orne Harbour | -64.61667 | -62.53333 |
| OE5-03 | 4/12/2024 | Antarctic Peninsula | D'Hainaut Island (Mikkelsen Harbour) | -63.9 | -60.78333 |
| OE5-04 | 10/12/2024 | South Shetland Islands | Pendulum (Deception Island) | -62.9333 | -60.6 |
| OE5-04 | 10/12/2024 | South Shetland Islands | Telefon Bay (Deception Island) | -62.93333 | -60.66667 |
| OE5-04 | 11/12/2024 | Antarctic Peninsula | Cuvertville Island | -64.68333 | -62.63333 |
| OE5-04 | 13/12/2024 | Antarctic Peninsula | D'Hainaut Island (Mikkelsen Harbour) | -63.9 | -60.78333 |
| OE5-04 | 14/12/2024 | Antarctic Peninsula | Gourdin | -63.2 | -57.3 |
| OE5-04 | 15/12/2024 | South Shetland Islands | Half Moon Island | -62.59 | -59.91 |
| OE5-04 | 15/12/2024 | South Shetland Islands | Yankee Harbour (Greenwich Island) | -59.78333 | -62.53333 |
| Argus | 14/12/2024 | South Shetland Islands | Ardley Island (King George Island) | -62.21308 | -58.93317 |
| Argus | 15/12/2024 | Antarctic Peninsula | Cierva Cove | -64.15 | -60.88333 |
| Argus | 16/12/2024 | Antarctic Peninsula | Dorian Bay | -64.81667 | -63.5 |
| Argus | 16/12/2024 | Antarctic Peninsula | Port Lockroy | -64.82528 | -63.49444 |
| Argus | 17/12/2024 | Antarctic Peninsula | Torgersen Island | -64.76667 | -64.08333 |
| Argus | 18/12/2024 | Antarctic Peninsula | Yalour Islands | -65.23333 | -64.16667 |
| Argus | 19/12/2024 | Antarctic Peninsula | Petermann Island | -65.16667 | -64.16667 |
| OE5-06 | 1/1/2025 | Antarctic Peninsula | Danco Island | -64.73333 | -62.61667 |
| OE5-06 | 2/1/2025 | Antarctic Peninsula | Cuvertville Island | -64.68333 | -62.63333 |
| OE5-06 | 2/1/2025 | Antarctic Peninsula | Portal Point | -64.5 | -61.76667 |
| OE5-06 | 3/1/2025 | Antarctic Peninsula | Palaver Point (Two Hummock Island) | -64.13333 | -61.7 |
| OE5-07 | 11/2/2025 | Antarctic Peninsula | Petermann Island | -65.16667 | -64.16667 |
| OE5-07 | 11/2/2025 | Antarctic Peninsula | Yalour Islands | -65.23333 | -64.16667 |
| OE5-07 | 12/2/2025 | Antarctic Peninsula | Danco Island | -64.73333 | -62.61667 |
| OE5-07 | 12/2/2025 | Antarctic Peninsula | Georges Point | -64.66667 | -62.66667 |
| OE5-07 | 13/2/2025 | Antarctic Peninsula | Portal Point | -64.5 | -61.76667 |

**Table S2.**
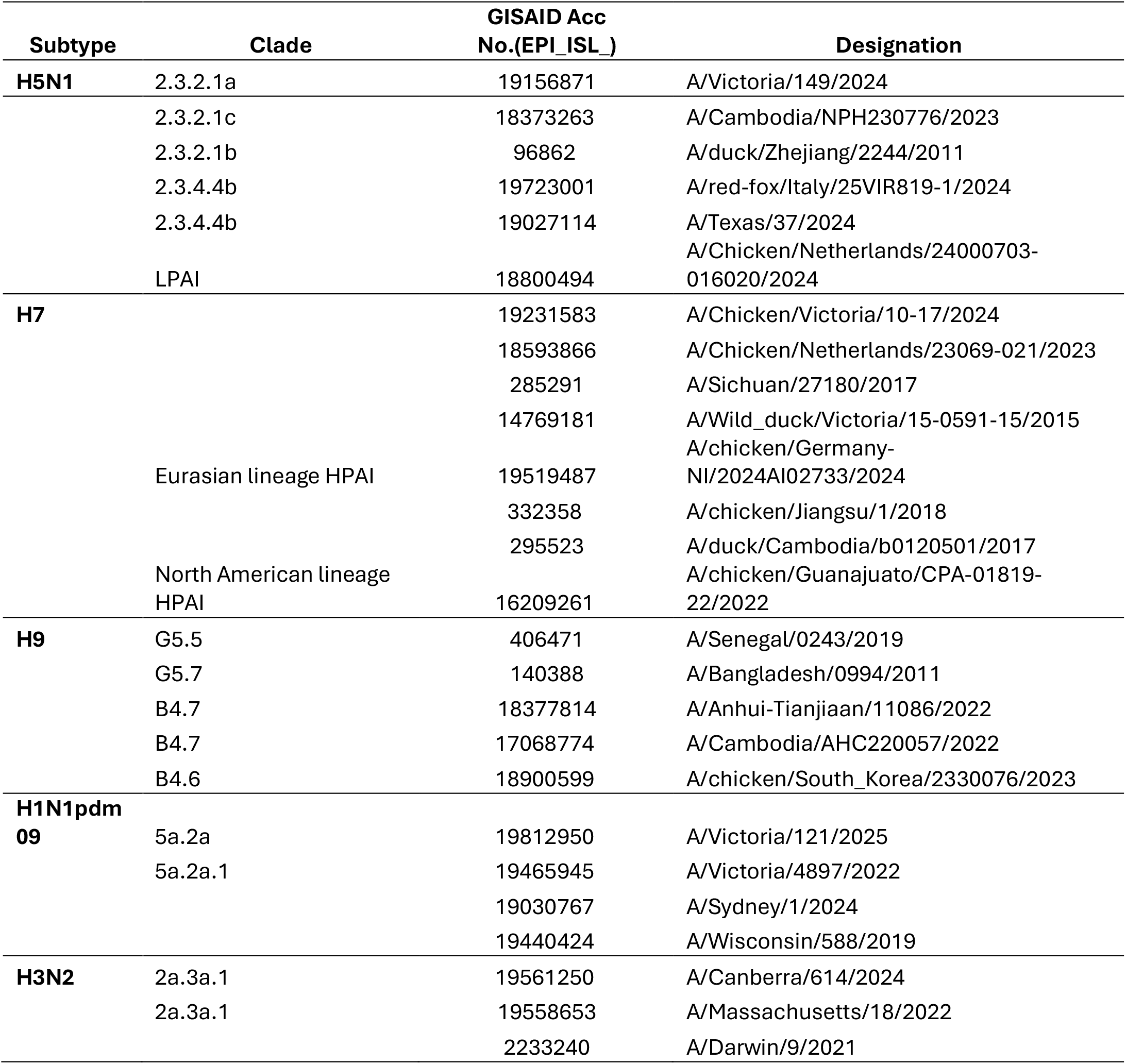
Influenza A genomes used for initial design of TWIST panel. The influenza B virus, respiratory syncytial viruses (RSV) and human metapneumovirus (hMPV) genomes incorporated are not shown.

**Figure S1.**
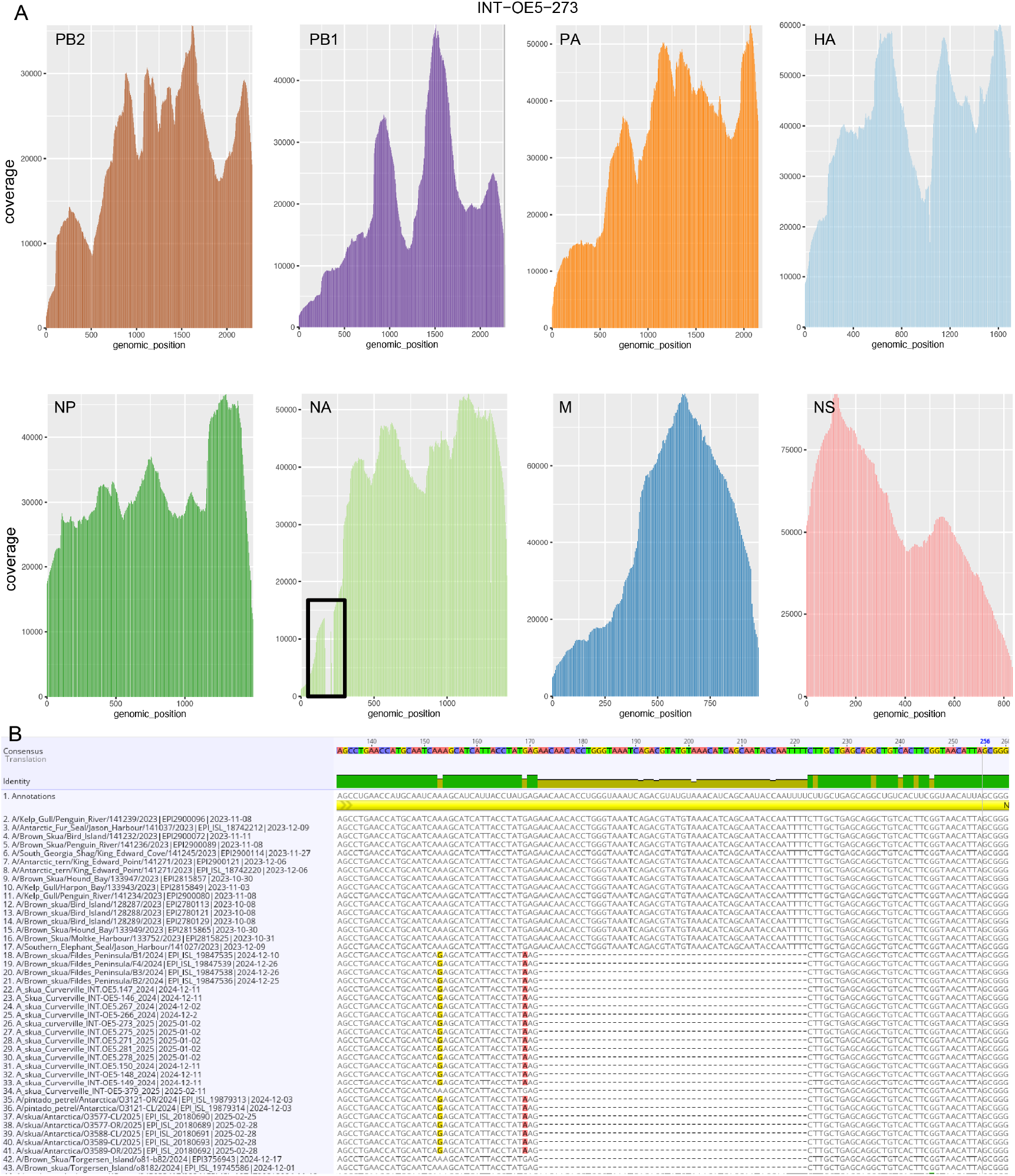
(A) Coverage map for a representative genome generated in this study, A/ Skua/Curverville/INT-OE5-273/2024(H5N1). Each panel comprises a segment. For the NA segment, across all genomes, a 51 bp detection was detected relative to the reference genomes. Mean coverage for all genomes is provided in Table S3 (B) Excerpt of alignment of the NA-N1 of sequences from the sub-Antarctic and Antarctica demonstrating this deletion to occur in genomes from [22,32,33], suggesting a *bona fide* deletion.

**Table S3. Ct values, coverage statistics, and accession numbers for positive samples from Skuas.**

See attached excel spreadsheet

**Table S4. Mutations of note identified using Flu Server.**

See attached excel spreadsheet

